# Trans-cellular tunnels induced by the fungal pathogen *Candida albicans* facilitate invasion through successive epithelial cells without host damage

**DOI:** 10.1101/2021.09.09.459426

**Authors:** Joy Lachat, Alice Pascault, Delphine Thibaut, Rémi Le Borgne, Jean-Marc Verbavatz, Allon Weiner

## Abstract

The opportunistic fungal pathogen *Candida albicans* is normally commensal, residing in the mucosa of most healthy individuals. In susceptible hosts, its filamentous hyphal form can invade epithelial layers leading to superficial or severe systemic infection. Invasion is mainly intracellular, though it causes no apparent damage to host cells. We investigated the invasive lifestyle of *C. albicans in vitro* using live-cell imaging and the damage-sensitive reporter galectin-3. Quantitative single cell analysis showed that invasion can result in host membrane breaching at different stages of invasion and cell death, or in traversal of host cells without membrane breaching. Membrane labelling and three-dimensional “volume” electron microscopy revealed that hyphae can traverse several host cells within trans-cellular tunnels that are progressively remodelled and may undergo ‘inflations’ linked to host glycogen stores, possibly during nutrient uptake. Thus, *C. albicans* invade epithelial tissues by either inflicting or avoiding host damage, the latter facilitated by trans-cellular tunnelling.

## Introduction

The human fungal pathogen *Candida albicans* (*C*.*a*) colonizes the oral, genital and intestinal mucosa of most healthy individuals and is part of the normal commensal flora (Gow *et al*., 2011). *C*.*a* often causes superficial infections such as oral or vaginal thrush, and in susceptible hosts can invade the gastrointestinal mucosa and enter the bloodstream, leading to a severe and life-threatening systemic infection (Pappas *et al*., 2018). *C*.*a* possesses a variety of virulence traits, allowing it to colonize within the microbiota when commensal, and invade host tissues during infection. *C*.*a’* ability to undergo a morphological switch, transitioning from round yeast cells to a filamentous hyphal form in response to various environmental cues, is considered to be its most important virulence trait, and has been strongly linked to its ability to invade and damage host tissue (Yang *et al*., 2014; Höfs, Mogavero and Hube, 2016).

Infection of epithelial layers proceeds through a series of sequential steps: attachment, initiation of hyphal growth, invasion and damage (Wächtler *et al*., 2011). Attachment to the host cell surface is mediated by various adhesins, such as Als3, which binds to E-cadherin receptor on epithelial cells, as well as Hwp1, Eap1 and Iff4 among others (Sheppard *et al*., 2004; Phan *et al*., 2007; Zhu and Filler, 2010).

Invasion is thought to be mainly an intracellular (transcellular) process, at least until the loosening of epithelial connections and barrier breakdown (Scherwitz, 1982; Yan, Yang and Tang, 2013; Goyer *et al*., 2016; Allert *et al*., 2018). Intracellular invasion is supported by two-dimensional electron microscopy (EM) studies as well as differential staining invasion assays, suggesting invasion originates predominantly at the apical surface of epithelial cells (Scherwitz, 1982; Zink *et al*., 1996; Zakikhany *et al*., 2007; Dalle *et al*., 2010; Wächtler *et al*., 2011).

While invasion begins around one or two hours after attachment, host cell damage, measured by LDH or ^51^Cr release assays, has been reported to occur at a later stage, usually after 6-24 hours, depending on the epithelial tissue (Zakikhany *et al*., 2007; Dalle *et al*., 2010; Sun *et al*., 2010; Böhringer *et al*., 2016). Indeed, the major cause of damage during *C*.*a* epithelial infection is thought not to be invasion *per se*, but other factors such as Candidalysin, a fungal peptide toxin secreted during infection (Moyes *et al*., 2016). Epithelial cells infected by a mutant deficient in secretion of Candidalysin are still invaded normally, but suffer no apparent damage (Moyes *et al*., 2016; Wilson, Naglik and Hube, 2016; Allert *et al*., 2018; Basmaciyan *et al*., 2019). How *C*.*a* intracellular invasion can proceed without damaging host cells is currently not well understood (Westman, Hube and Fairn, 2019). While hyphal extension and the resulting invagination of the host plasma membrane into the so-called ‘invasion pocket’ can account for at least a portion of intracellular invasion, further hyphal extension within the host, and even more so into neighbouring host cells, is expected to lead to eventual host membrane breaching and damage due to increased membrane stretching (Mogavero *et al*., 2021; Swidergall *et al*., 2021). Current models hypothesize the presence of a non-damaging intracellular invasion route, though the mechanism underlying such a route has not been described (Allert *et al*., 2018; Basmaciyan *et al*., 2019).

Two mechanisms of *C*.*a* invasion are known: induced endocytosis and active penetration (Richardson, Ho and Naglik, 2018; Basmaciyan *et al*., 2019). Induced endocytosis is considered a “host-driven” process, whereby invasins expressed on the surface of *C*.*a* hyphae induce host cells to produce pseudopods that engulf the pathogen and pull it inside the cell, a process analogous to the “trigger” mechanism employed by several invasive bacteria such as *Shigella* and *Salmonella* (Drago *et al*., 2000; Cossart and Sansonetti, 2004; Park *et al*., 2005; Zakikhany *et al*., 2007; Dalle *et al*., 2010). In epithelial cells, the invasins Als3 or Ssa1 activate an E-cadherin mediated, clathrin-dependent pathway, though an E-cadherin-independent pathway has been suggested as well (Phan *et al*., 2007; Moreno-Ruiz *et al*., 2009; Sun *et al*., 2010; Liu and Filler, 2011; Wächtler *et al*., 2012). Active penetration is considered a “fungal-driven” process that relies on a combination of physical forces exerted by extending hyphae and secreted fungal factors such as aspartyl proteinases (Saps), though still relatively little information about this mechanism exists (Julian R. Naglik *et al*., 2008; Naglik *et al*., 2011). The actin polymerization inhibitor Cytochalasin D has been reported to disrupt invasion by induced endocytosis but not by active penetration, marking a key distinction between these two mechanisms (Park *et al*., 2005; Dalle *et al*., 2010; Wächtler *et al*., 2012). While oral, vaginal and microfold (M) cells are invaded via both mechanisms, enterocytes are invaded only via active penetration, though the underlying reason for this cell type dependency is unclear (Dalle *et al*., 2010; Albac *et al*., 2016; Swidergall and Filler, 2017; Zhang *et al*., 2018). On the host side the response to invasion remains poorly described, though actin, clathrin, dynamin and cortactin have been shown to be recruited to the site of invasion and to have a functional role in *C*.*a* internalization in HEK293 and JEG-3 cells (Tsarfaty *et al*., 2000; Moreno-Ruiz *et al*., 2009). The small GTPases Rac1, Cdc42 and Rho were also shown to be co-localized with actin at the site of invasion, though their precise function in this context is not known (Atre *et al*., 2009).

In general, two strategies can be employed by invasive pathogens to enter host cells: breaching of host membranes and entry into the host cytosol or residence within a membrane-bound compartment. This distinction between cellular “niches” has been especially useful for describing the intracellular lifestyles of invasive bacteria which include: membrane breaching pathogens like *Shigella, Rickettsia* and *Listeria*; pathogens that reside in membrane-bound compartments like *Brucella, Legionella* and *Chlamydia;* and pathogens that can adapt both lifestyles like *Salmonella* and *Mycobacterium tuberculosis* (Cossart and Sansonetti, 2004; Simeone *et al*., 2012; Cossart and Helenius, 2014; Fredlund *et al*., 2018). The fungal pathogen *Cryptococcus neoformans* resides within a membrane-bound compartment in macrophages, where it can induce non-lytic expulsion through fusion of the phagosome and plasma membranes (Johnston and May, 2010). In contrast, the invasive lifestyle of *C*.*a* is still poorly understood, with limited information derived mostly from end-point assays and two-dimensional EM regarding the host cellular niches encountered during epithelial invasion (Scherwitz, 1982; Zink *et al*., 1996; Park *et al*., 2005; Zakikhany *et al*., 2007; Dalle *et al*., 2010). Indeed, it is currently unclear if and when host membranes are breached during *C*.*a* invasion; a key aspect in understanding this pathogen’s invasive lifestyle.

## Results

### New experimental pipeline combines the galectin-3 damage sensitive reporter and live-cell imaging to study *C*.*a* invasion

We developed a new experimental pipeline to study membrane breaching events during *C*.*a* epithelial cell invasion at the single cell level using the damage sensitive reporter galectin-3 combined with multi-dimensional live-cell imaging and other cell markers. Galectin-3 is a small soluble protein localized in the cytosol, which has affinity to β-galactose-containing carbohydrates. These moieties are typically found in vesicles and at the outer leaflet of the plasma membrane, but not in the cytosol nor the nucleus (Paz *et al*., 2010). By expressing galectin-3 linked to a fluorescent reporter, the precise location and timing of membrane breaching events can be tracked in real-time, as upon damage fluorescent galectin-3 flows from the cytosol, through the ruptured membrane and onto the cell surface, where it binds to moieties not normally accessible to it (Paz *et al*., 2010). Galectin-3 has been used to study the lifestyles of invasive bacteria such as *Shigella flexneri* and *Salmonella typhimurium* (Paz *et al*., 2010; Ray *et al*., 2010; Mellouk *et al*., 2014; Weiner *et al*., 2016; Fredlund *et al*., 2018), as well as *C*.*a* escape from macrophages (Westman *et al*., 2020). In short, the experimental pipeline consists of the following steps (see **Figure 1A**): (1) *C*.*a* are cultured to log phase, diluted to a multiplicity of infection (MOI) of 0.1. In parallel, epithelial cells stably expressing eGFP-Gal-3 (from here on referred to as ‘Gal-3’) are cultured to confluency followed by staining with CellMask (CM), a membrane stain suitable for live-cell imaging which labels the entire host plasma membrane and endocytic compartment. Next, *C*.*a* are added above the epithelial layer, followed by live-cell imaging of multi-channel z-stacks acquired every 10 minutes for 7 hours.

**Figure 1.**
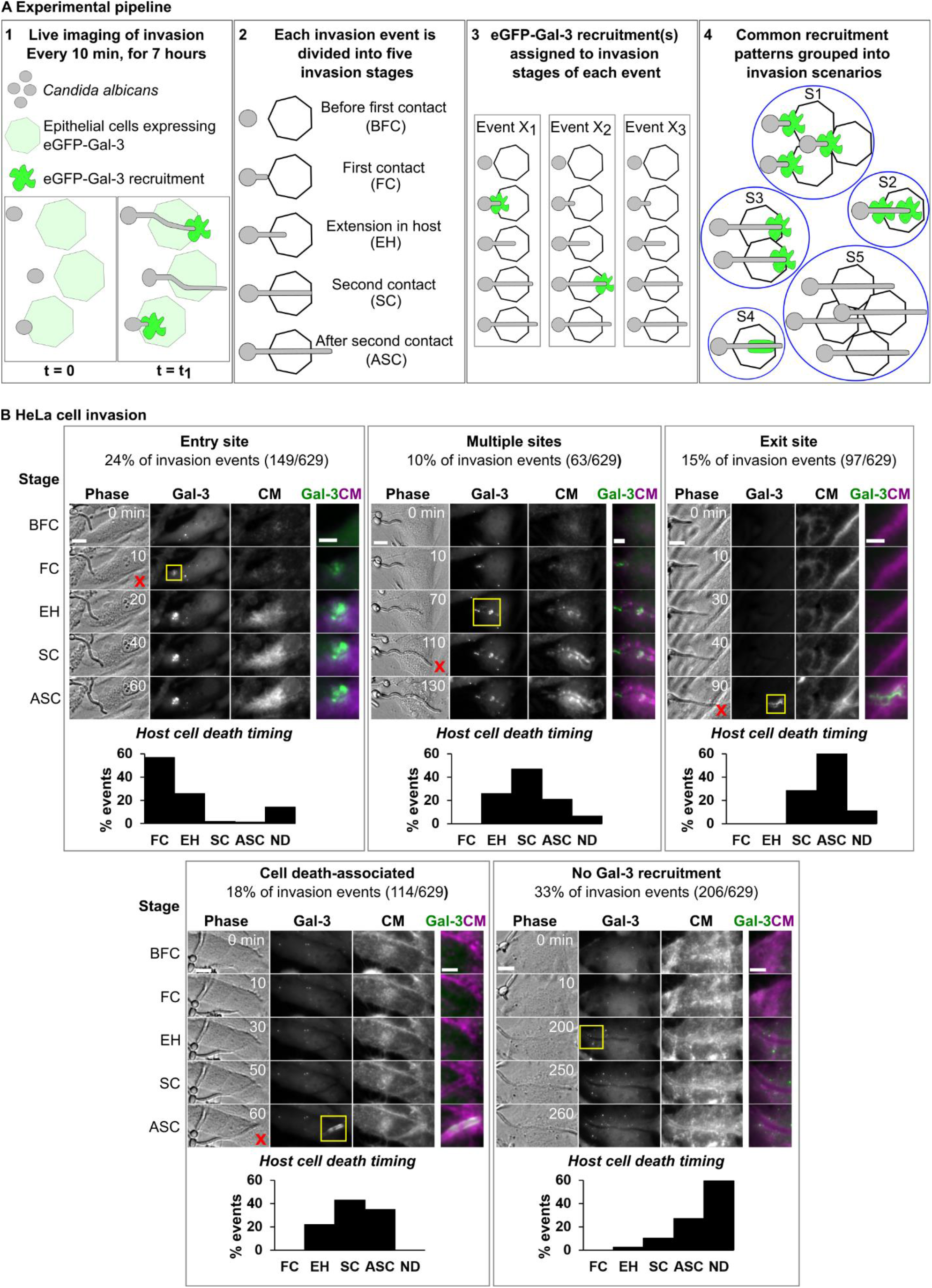
*C*.*a* epithelial cell invasion using eGFP-Galectin-3 damage sensitive reporter and live-cell imaging. The experimental pipeline (A) and results for HeLa invasion (B) are presented. (A1) *C*.*a* invasion of epithelial cells stably expressing eGFP-Galectin-3 is studied using live-cell imaging. Localized high-intensity eGFP-Galectin-3 recruitments recorded are indicative of host membrane damage. (A2) Each invasion event, defined as a single interaction between a *C*.*a* hypha and a host cell over time, is sub-divided into five invasion stages: before first contact with host plasma membrane (BFC), first contact with host plasma membrane (FC), hypha extension within the host (EH), second contact with the host plasma membrane (SC), after second contact with the host plasma membrane (ASC). (A3) For every invasion (X_1_, X_2_ ..), eGFP-Galectin-3 recruitments are assigned to the stage in which they occurred. (A4) All invasion events sharing common eGFP-Galectin-3 recruitment patterns are grouped into invasion scenarios (S_1_, S_2_ ..). (B) *C*.*a* HeLa invasion scenarios (n = 629 invasion events). For each scenario a representative invasion event is presented, divided into the five invasion stages described in (A). For each stage a single section is presented in phase, Gal-3, and CM channels, as well as an inset (yellow frame) containing a composite view of the Gal-3 (green) and CM (magenta) channels. Cell death (if it occurred) is marked with a red **x** in the corresponding invasion stage. The distribution of cell death timing as a function of the invasion stage is presented for each scenario. ND stands for no cell death during invasion. Scale bars are 10 µm or 5 µm in insets.

(2) Post-acquisition, every *invasion event*, defined as an interaction between a single *C*.*a* hypha and a single host cell, is identified and analysed individually. Each event is sub-divided into five *invasion stages*: 1. Before first contact with the host plasma membrane (BFC), 2. First contact with the host plasma membrane (FC), 3. Extension within the host cell (EH), 4. Second contact with host plasma membrane (SC), 5. After second contact with the host plasma membrane (ASC).

(3) Observed Gal-3 recruitments are assigned to their corresponding invasion stage.

(4) All invasion events presenting common stage-dependant Gal-3 recruitment patterns are grouped into *invasion scenarios* that represent the different ways in which host membrane damage (or lack thereof) is manifested during *C*.*a* invasion into a specific cell line. During the analysis, the localization patterns of CM and host cell death timing (determined by membrane blebbing and condensed nuclei observed in phase contrast (Morgan *et al*., 2002)) are also recorded and assigned to their corresponding invasion stage for every invasion event. However, only the Gal-3 signal is used to define the different invasion scenarios (for full pipeline details see **Materials and Methods**).

### Invasion into HeLa cells occurs via five invasion scenarios

We first studied *C*.*a* invasion into HeLa cells (from here on referred to as HeLa) expressing Gal-3, as this non-polar cell line has been previously used to characterize in detail host membrane damage and the invasive lifestyles of *Shigella* and *Salmonella* (Mellouk *et al*., 2014; Fredlund *et al*., 2018). HeLa have also been used to study *C*.*a* invasion, with both induced endocytosis and active penetration implicated as the mechanisms of invasion (Drago *et al*., 2000; Moreno-Ruiz *et al*., 2009; Wächtler *et al*., 2012; Maza *et al*., 2017). Wild-type *C*.*a* strain BWP17 (WT strain) (Bassilana, Hopkins and Arkowitz, 2005) invasion into HeLa expressing Gal-3 was studied using the experimental pipeline described above. Overall, we studied 629 invasion events in six statistically comparable experiments.

Analysis of invasion events revealed five distinct invasion scenarios (see **Figure 1B, Movies 1-5**): ‘Entry site’ scenario: Gal-3 is recruited at the site of contact between hyphae and the host plasma membrane during FC (24% of events); ‘Multiple sites’ scenario: Gal-3 is recruited at multiple sites during FC and EH (10% of events); ‘Exit site’ scenario: Gal-3 is recruited at the second contact with the host plasma membrane during SC or ASC (15% of events); ‘Cell death-associated’ scenario: Gal-3 is recruited in an elongated pattern along the hypha trajectory. This recruitment pattern always coincided with host cell death, as revealed by cell death timing data (see details below) (18% of events); ‘No Gal-3 recruitment’ scenario: no Gal-3 recruitment is observed during invasion (33% of events). Overall, we observe that ‘damaging scenarios’ exhibiting Gal-3 recruitments constitute 67% of invasion events, with the ‘non-damaging’ ‘no Gal-3 recruitment’ scenario constituting the rest.

We calculated the host cell traversal time, defined as the time it takes for a hypha to extend from the FC stage to the SC stage, if no host cell death occurs. In HeLa, traversal time averages 70 minutes ± 38 min independently of the invasion scenario, suggesting host membrane breaching does not impact the overall dynamics of invasion (see **Figure S1A**). In order to confirm that Gal-3 recruitments were associated with internalized hyphae extending inside host cells, we used a fixed differential staining assay adapted from previous invasion studies (Park *et al*., 2005; Dalle *et al*., 2010) (see **Figure S2**). Of note, while invading hyphae were often observed extending in proximity to or alongside host cell nuclei, *C*.*a* invasion into the nucleus was never observed.

The distribution of host cell deaths as a function of invasion stage was calculated for every invasion scenario (see **Figure 1B)**. During ‘damaging scenarios’ host cell death was observed in 92% of events, compared to 41% of events in the ‘no Gal-3 recruitment’ scenario, with two thirds of deaths in the latter scenario occurring in the ASC stage. The elongated Gal-3 recruitment pattern observed in the ‘cell death-associated’ scenario always coincided with host cell death, within the temporal resolution of the experiment. This recruitment pattern is likely a result of general host cell membrane permeabilization allowing Gal-3 to bind membranes surrounding the hyphae’s path (see below). Taken together, host membrane breaching during HeLa invasion is most often associated with host cell death, while invasion without host membrane breaching is most often associated with host cell survival.

Analysis of the host plasma membrane and endocytic compartment marker CM revealed a faint but clear uniform labelling of host cell membranes along the path of invasion during the ‘exit site’, ‘cell death-associated’ and the ‘no Gal-3 recruitment’ scenarios. In contrast, in the ‘entry site’ scenario, CM labelling was found only at the site of membrane breaching, and in the ‘multiple sites’ scenario, CM labelling was in the form of discontinuous puncta and patches along the path of invasion.

Next, we used our experimental pipeline to study the three-dimensional dynamics of CM recruitment in detail. In one striking example, a single HeLa cell was invaded by two hyphae, each via a different invasion scenario (see **Figure 2, Movie 6**). Invasion by the first hypha (I) started with a Gal-3 recruitment during the FC stage, followed by smaller Gal-3 recruitments during the EH stage, indicating a ‘multiple sites’ invasion scenario. Around this hypha, only localized, discontinuous CM recruitment was observed along the path of invasion. Invasion by the second hypha (II) did not trigger Gal-3 recruitment until the ASC stage. This hypha was tightly surrounded by a continuous CM label throughout invasion. During the SC stage, a clear host plasma membrane outward deformation was observed, followed by a ring-like Gal-3 recruitment during the ASC stage, indicating an ‘exit site’ invasion scenario. A cross-section view of the CM signal through both hyphae shows the absence of CM around hypha I, and a circular CM organization around hypha II. These CM recruitment patterns were representative of all ‘multiple sites’ and ‘exit site’ scenarios studied. Overall, we observe that CM organization during HeLa invasion is dependent on the invasion scenario, with a continuous circular organization around invading hyphae in the ‘no Gal-3 recruitment’, ‘exit site’ and ‘cell-death associated’ scenarios, a patchy and discontinuous organization along the path of invasion in the ‘multiple sites’ scenario, and a localized recruitment at the site of membrane breaching in the ‘entry site’ scenario (see **Figure S3**).

**Figure 2.**
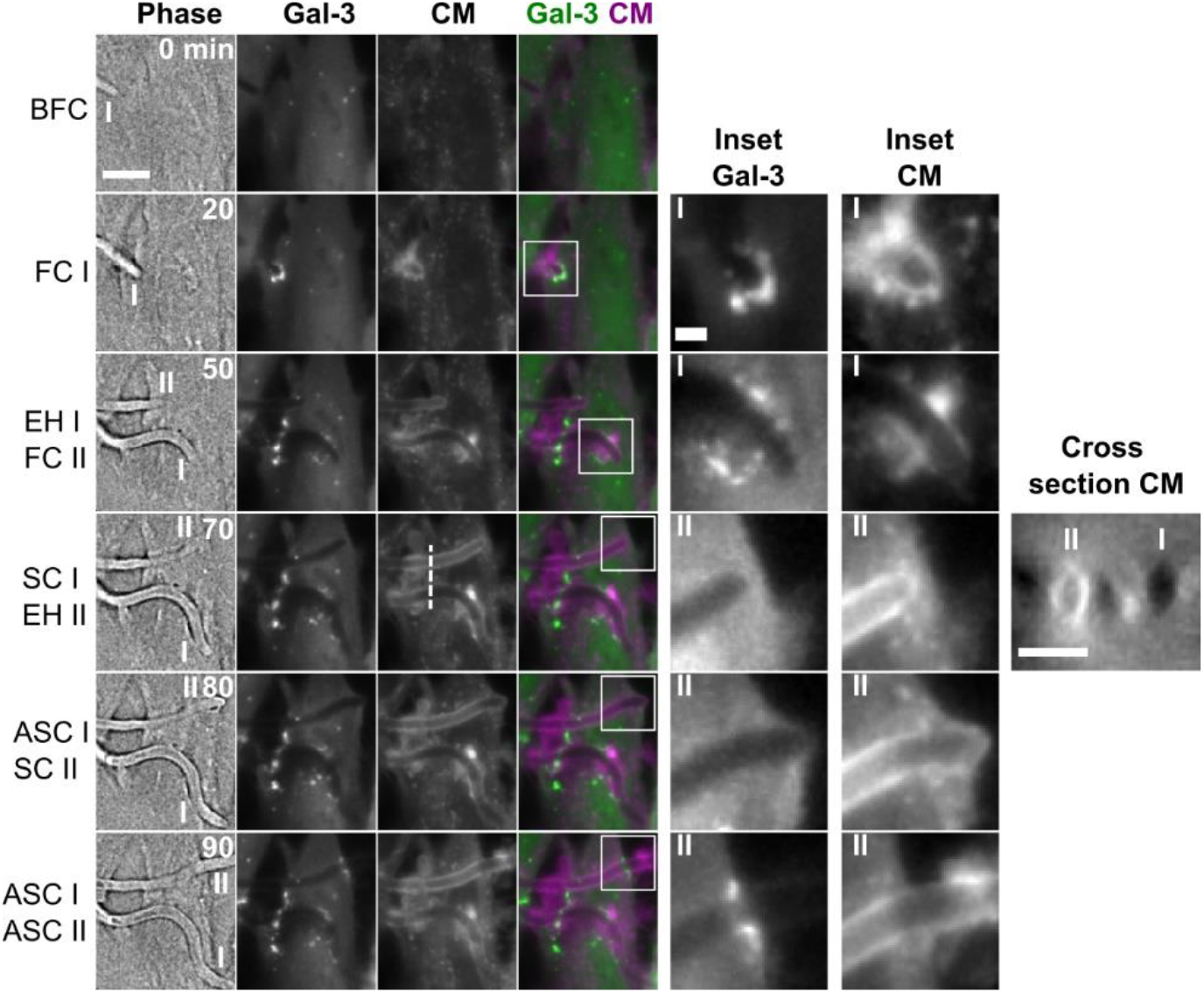
High-resolution live-cell imaging of two *C*.*a* hyphae invading a single HeLa cell via a ‘multiple sites’ scenario and an ‘exit site’ scenario. Six time points are presented, with the invasion stages defined in Figure 1 corresponding to either filament I (‘multiple sites’ scenario) or II (‘exit site’ scenario). For each time point a phase image is presented together with a maximum intensity projection of three z-sections in the Gal-3 and CellMask (CM) channels, and a composite image showing Gal-3 (green) and CM (magenta) together. Two insets (Gal-3 and CM) are presented for each timepoint, corresponding to filament I or II. A cross-section in the CM channel through both filaments (dotted line) is presented at time point 70 min. A circular CM organization is observed around filament II but not filament I. No host cell death was observed during the course of live-cell imaging. Scale bars are 10 µm, 2 µm (inset) and 5 µm (cross-section). For other invasion scenarios see **Figure S3**.

### Invasion into Caco-2 cells occurs via a single non-damaging invasion scenario

The Caco-2 cell line is a widely used model for enterocytes and is of particular relevance to *C*.*a* invasion, as the gut is considered to be the reservoir for severe systemic infection (Nucci and Anaissie, 2001; Grosheva *et al*., 2020). Several studies have indicated that *C*.*a* invasion into polarized Caco-2 cells (from here on referred to as Caco-2) occurs only via active penetration (Dalle *et al*., 2010; Albac *et al*., 2016; Goyer *et al*., 2016). No host damage (measured by LDH release) was detected in the first hours of Caco-2 infection, despite extensive invasion already taking place, suggesting that early infection in this cell line is non-damaging (Dalle *et al*., 2010; Böhringer *et al*., 2016; Allert *et al*., 2018).

Invasion into the Caco-2 clone C2BBe1 stably expressing Gal-3 was studied in 153 invasion events in three independent experiments, using the same experimental pipeline used above. Caco-2 polarization was confirmed by the presence of microvilli observed by EM (see below) (Rey *et al*., 2020). In contrast to HeLa invasion, invasion into Caco-2 occurred via a single invasion scenario in which no detectable Gal-3 recruitment and no host cell death took place during live-cell imaging (see **Figure 3A, Movie 7**). In all invasion events, continuous CM labelling along the entire path of invading hyphae was observed. Interestingly, when several host cells were invaded in sequence, CM labelling appeared to be continuous, with no detectable gaps during the transition between cells (see **Figure 3A and 3B, Movie 8**). Cross-section views revealed a circular CM organization around invading hyphae, which appeared identical to the CM labelling observed during three of the HeLa invasion scenarios (‘no Gal-3 recruitment’, ‘exit site’ and ‘cell-death associated’). This circular organization could also be detected in a head-on view during early invasion when hyphae were nearly aligned along the microscopical z-axis (see **Figure 3C**).

**Figure 3.**
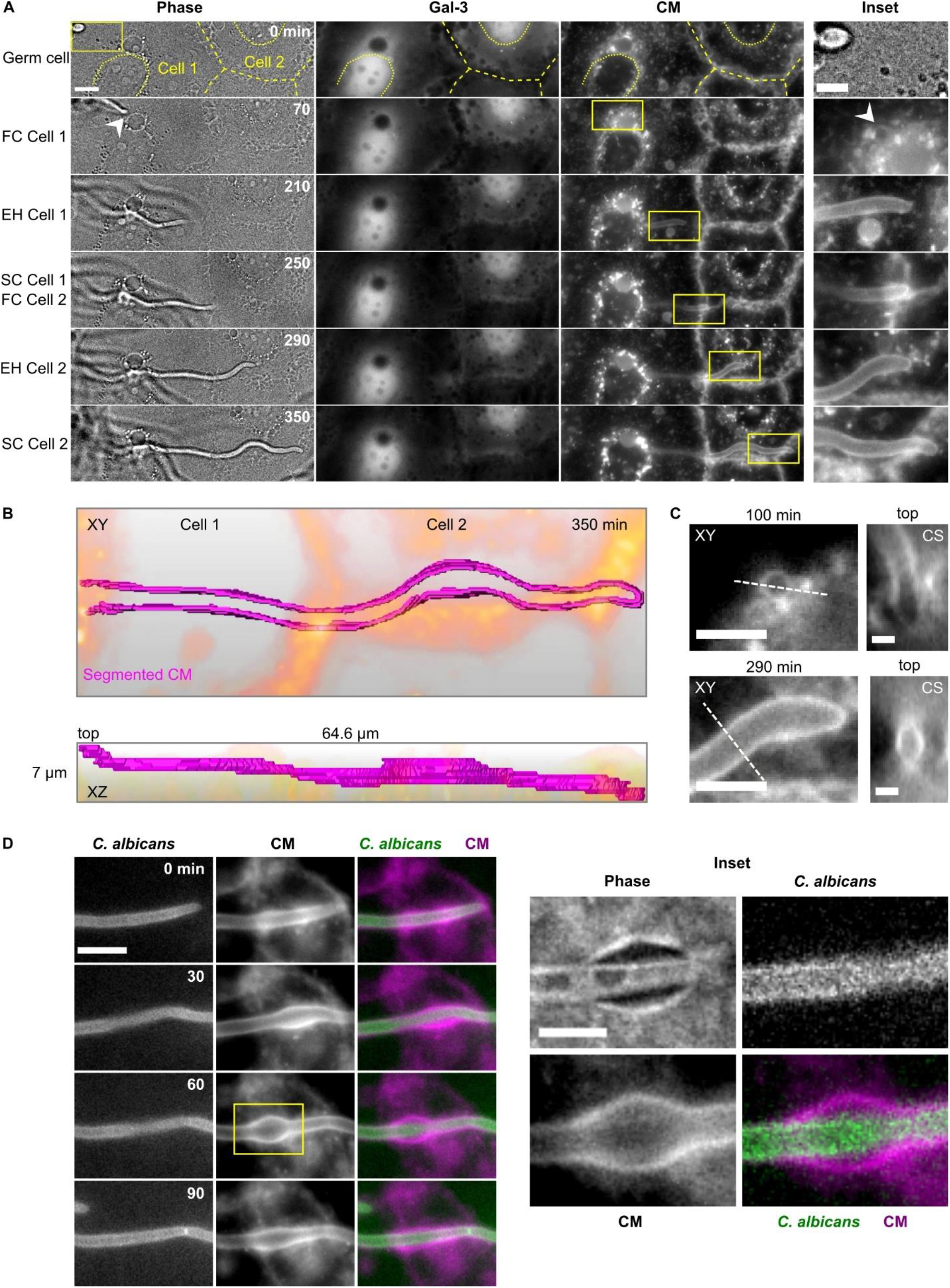
Invasion into Caco-2. (A) Caco-2 expressing Gal-3 are invaded by a single non-damaging invasion scenario without Gal-3 recruitment or cell death. A representative live-cell imaging acquisition of invasions through two successive cells (cell 1 and cell 2) is presented. The six time points correspond to the invasion stages defined in Figure Phase (single z slice), Gal-3 and CM (maximum intensity projection of 7 slices) channels, together with an inset (yellow frame) are presented for each time point. The outlines of the two invaded Caco-2 and their nuclei are highlighted with dashed and dotted yellow lines respectively. The entry point into the Caco-2 layer is marked by an arrowhead. (B) Three-dimensional representation of CM labelling (magenta) presented in (A) within the entire acquisition volume during a single time point (350 min). Epithelial cell outline is in orange. A top view (XY) and side view (XZ) are presented together with the volume dimensions. (C) CM channel insets of two time points from the data presented in (A) are presented together with cross-section (CS) views derived from the dashed lines. (D) Live-cell imaging of infection of Caco-2 not expressing Gal-3 with a *C*.*a* strain expressing GFPγ. An average intensity projection of 5 slices is presented. A local transient ‘inflation’ is observed in the CM channel (left). An inset showing a clear separation between the CM labelling and *C*.*a* when the ‘inflation’ is at its maximum size is presented. Scale bars are 10 µm in (A) and (D), 5 µm in (A) Inset, (C) XY views and (D) inset, and 2 µm in (C) CS views.

The host cell traversal time during Caco-2 invasion was 97 ± 54 minutes (see **Figure S1B**) and hyphae internalization was confirmed using a fixed differential staining assay (see **Figure S2**). As in HeLa, nuclear invasion was never observed in Caco-2.

In 25% of invasion events (n = 38/153) we observed a localized and transient expansion of the CM labelling surrounding invading hyphae. In order to visualize these membrane ‘inflations’ in detail, we performed invasion experiments into Caco-2 (not expressing Gal-3), using a *C*.*a* strain expressing GFPγ at the plasma membrane, allowing clear distinction between the CM signal and the hypha outline (see **Figure 3D, Movie 9**). Each inflation originated from the typical tight CM labelling around extending hyphae, then grew bigger over time until a clear separation between the CM signal and the hypha was detectable (see **Figure 3D inset**), followed by a return to a tight CM labelling around the extending hypha. Inflations were observed along the entire path of invasion through Caco-2, and were never observed during HeLa invasion.

### Cytochalasin D treatment does not alter HeLa or Caco-2 invasion

As induced endocytosis, but not active penetration, is inhibited by the actin polymerization inhibitor cytochalasin D, we examined the effect of cytochalasin D treatment on HeLa and Caco-2 invasion using our experimental live-cell imaging Gal-3 pipeline. We found that in both HeLa (see **Figure 4A**) and Caco-2 invasion (see **Figure 4B**), treatment did not significantly alter the number of invasion events. The distribution of HeLa invasion scenarios was also not significantly altered, while invasion into treated Caco-2 occurred via a single non-damaging invasion scenario as in untreated cells. These results demonstrate that invasion in these cell types does not occur via induced endocytosis, in agreement with previously published results relating to Caco-2 (Dalle *et al*., 2010).

**Figure 4.**
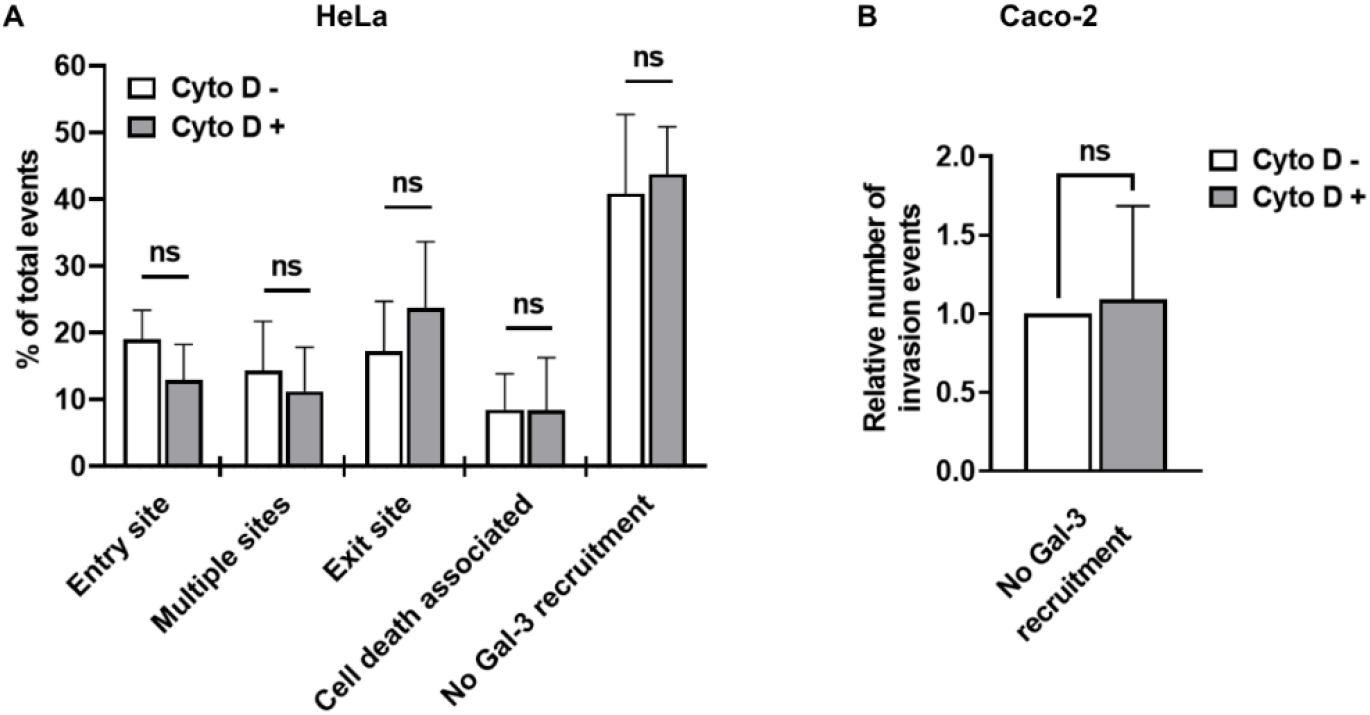
Cytochalasin D treatment does not significantly alter the number of invasion events in HeLa or Caco-2. (A) HeLa. n = 618 invasion events. (B) Caco-2, n = 261 invasion events.

### Serial block face-scanning electron microscopy (SBF-SEM) of Caco-2 invasion provides a three-dimensional nano-scale view of hyphae invading successive host cells

Understanding precisely how *C*.*a* invade host cells without damage requires imaging at a resolution below the limit of standard light microscopy. Moreover, studying cellular structures that extend over long cellular distances and even several cells requires three-dimensional nano-scale imaging of large sample volumes, a combination difficult to obtain using traditional EM approaches (Weiner and Enninga, 2019). We therefore studied invasion using serial block face-scanning electron microscopy (SBF-SEM), an emerging “volume” EM technique capable of imaging entire *C*.*a* filaments invading multiple host cells within a single 3D data set at nano-scale resolution (Peddie and Collinson, 2014; Smith and Starborg, 2019) (see **Figure 5**). As invasion into Caco-2 occurs via a single non-damaging invasion scenario, we targeted this cell line for SBF-SEM. Caco-2 were infected by the WT *C*.*a* for 6 hours, followed by EM sample preparation, which resulted in excellent sample contrast for both *C*.*a* internal structures like the Spitzenkörper and for membranes and organelles of the host (Weiner *et al*., 2019) (see **Materials and Methods**). Host cells presented clear microvilli at their apical surface, indicating monolayer polarization. Data sets were acquired in three different sessions, with a resolution of 10 nm in the acquisition plane and an axial resolution of 100 nm. Overall 11 volumes were acquired containing 10 invasion events in which the entire invading hypha and surrounding environment were captured, and 30 invasions events in which only a partial segment of an invading hypha was captured. A representative data set containing a *C*.*a* filament invading three host cells in succession is presented (see **Figure 5A, Movie 10**).

**Figure 5.**
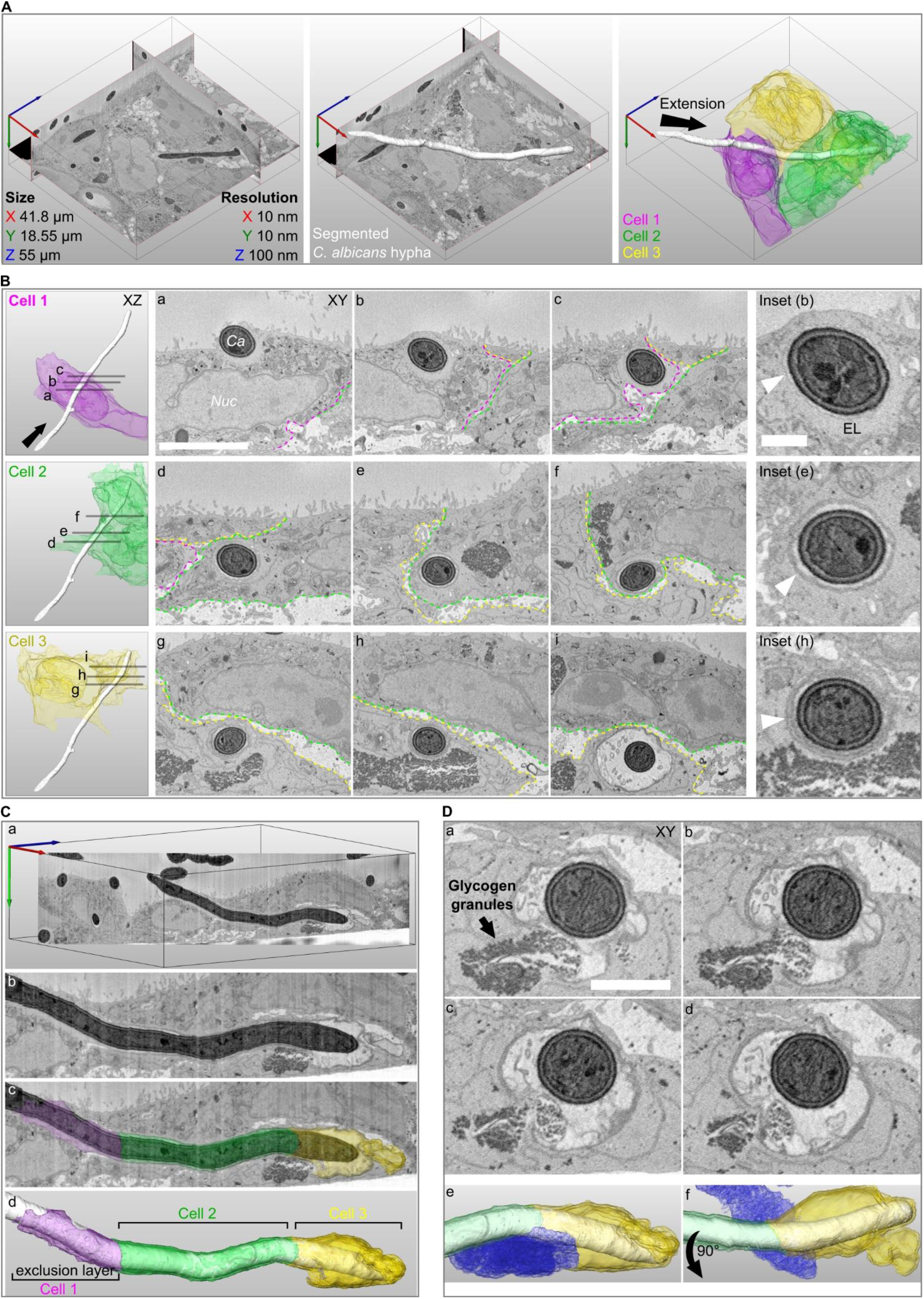
Serial block face-scanning electron microscopy (SBF-SEM) of Caco-2 invasion. (A) A representative SBF-SEM volume is presented together with data set size and resolution in each axis (left panel). A *C*.*a* hypha in the process of invasion is visualized in three-dimensions (white, middle panel). Three invaded host cells are colour coded according to the sequence of invasion, cell 1 (magenta), cell 2 (green), cell 3 (yellow). The direction of filament extension is noted (arrow, right panel). (B) The organization of host membranes around the hypha is presented for each invaded cell with three cross-section views (Cell 1- a, b, c; Cell 2- d, e, f; Cell 3- g, h, i). The direction of filament extension is noted (arrow). The borders between cells are annotated according to the colours in (A) and the hypha (Ca) and the host nucleus (Nuc) are noted in (a). An inset from the middle cross-section in each cell is presented with arrowheads pointing to host membranes surrounding the hypha. The hypha in cell 1 is tightly enveloped by a thin membrane which in turn is surrounded by an ‘exclusion layer’ (EL) devoid of cellular structures (a, b, c). In cell 2, a thicker membrane enveloping a narrow lumen and the hypha is observed (d, e, f). In cell 3, two membranes surrounding a narrow lumen and the hypha are observed in (g, h). An ‘inflation’ containing multiple vesicles is observed in (i). (C) A cross-section through the hypha long-axis is presented, showing the continuous membrane structures across three cells (a, b). The ‘exclusion layer’ and membranes surrounding the hypha are visualized according to the invaded cell sequence (c, d). (D) A sequence of four sequential sections showing direct contact between the ‘inflation’ lumen and host glycogen granules (a-d). Glycogen granules are noted in (a). A visualization presented in the same orientation as in (C) shows the contact between the host glycogen store (blue) and the ‘inflation’ in cell 3 (e). A 90-degree rotated view is presented in (f). Scale bars are *5* µm in B, 1 µm in B inset and 2 µm in D.

### Invading filaments are enveloped by host membranes in a tunnel structure progressively remodelled with each invaded cell

In all invasion events examined, invading hyphae were fully enveloped by host membranes on all sides (except at contact points with glycogen stores (see below)) so that hyphae were always found within a continuous tunnel structure that could extend through successively invaded host cells. In contrast, non-invading filaments observed at the Caco-2 monolayer surface were not associated with any external membranes. Careful examination of the data revealed that tunnel organization is linked to the sequence in which host cells were invaded (see **Figure 5B, Movie 11**). Invasion into the first host cell in sequence (cell 1) by a filament originating from the surface of the Caco-2 monolayer, was characterized by a tight enveloping of the invading filament by a single host membrane, which in turn was surrounded by an ‘exclusion layer’, that contained no host cellular compartments or organelles (see **Figure 5Ba and 5Bb**). Based on previously published work demonstrating localized actin recruitment at the site of *C*.*a* invasion into Caco-2, and ‘exclusion layers’ composed of actin seen during *S. flexneri* invasion, this ‘exclusion layer’ likely represents a region of actin enrichment (Dalle *et al*., 2010; Weiner *et al*., 2016). This ‘exclusion layer’ became progressively thinner as the filament extended through cell 1, eventually being enveloped again by the plasma membrane upon exit from cell 1 (see **Figure 5Bc**). Invasion into the second host cell in the sequence (cell 2) started with the enveloping of the structures surrounding the filament in cell 1 by the plasma membrane of cell 2 (see **Figure 5Bd**), forming a thicker, clearly visible membrane structure surrounding the hypha further into cell 2 (see **Figure 5Be**). This membrane structure persisted throughout cell 2 until the exit from the cell, where another enveloping by the plasma membrane of cell 2 took place (see **Figure 5Bf**). The transition between cell 2 and the third cell in the sequence (cell 3) appeared to occur in an identical manner, with an enveloping of the membrane structures seen in cell 2 (already containing structures from cell 1) by the plasma membrane of cell 3, so that two clearly distinguishable membrane layers (each likely containing multiple plasma membrane sheets, see **Discussion**) could be observed (see **Figure 5Bg and 5Bh**). Taken together, we observe a progressive remodelling of the tunnel structure surrounding invading filaments which occurs in correspondence to the sequence of invaded cells (see **Figure 5C, Movie 12** and model in **Figure 6**). This accumulation of membranes can also be clearly observed in the multi-layer membrane structures surrounding filaments that have invaded several cells (see **Figure S4**). Of note, the above sequence of progressive tunnel remodelling, including the presence of an exclusion layer in cell 1, was observed in every invasion event examined by SBF-SEM (n = 10 full tunnels and 30 partial tunnels).

**Figure 6.**
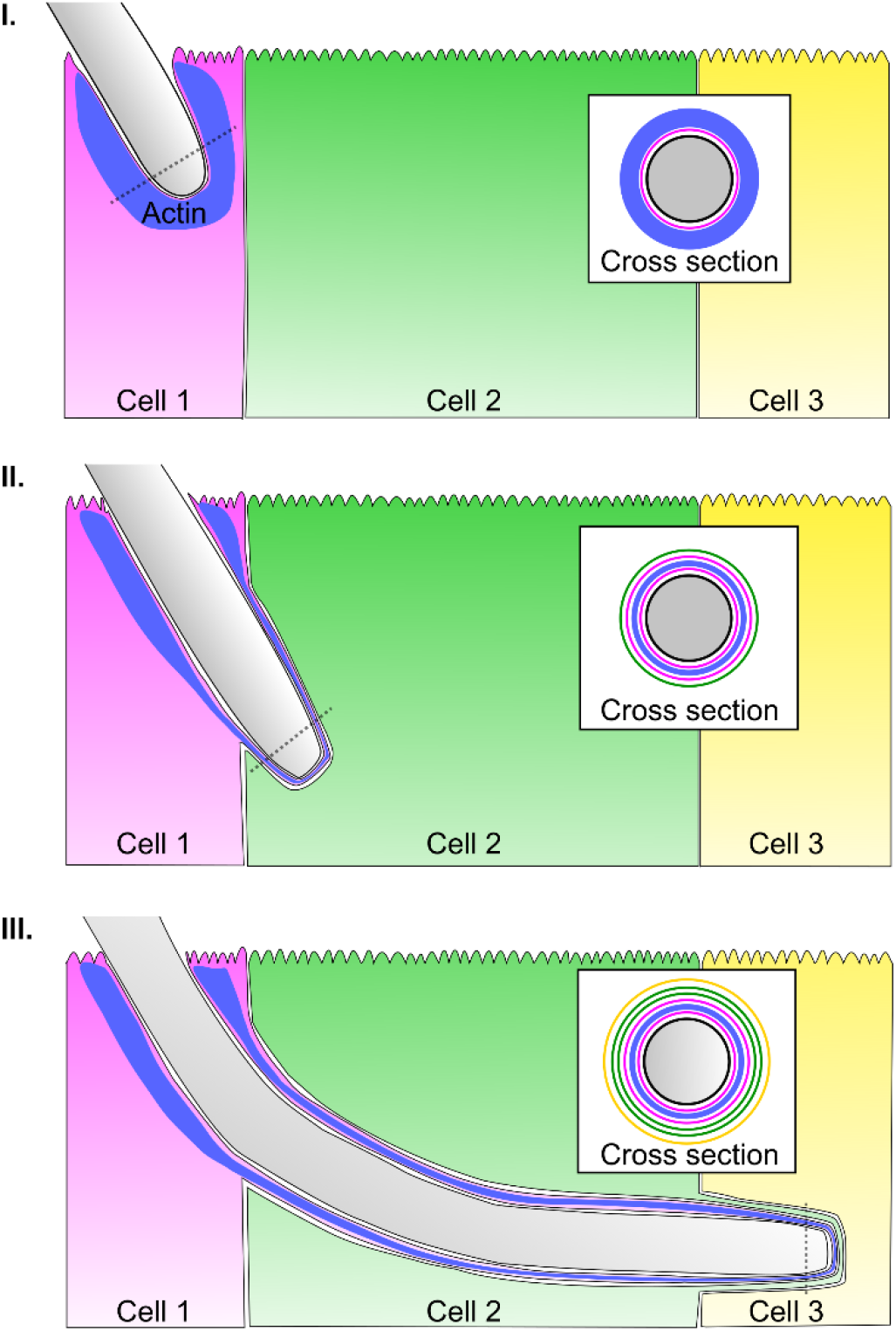
Model of *C*.*a* trans-cellular tunnelling (CaTCT). I. Invasion into the first cell in the invasion sequence (cell 1) is characterized by tight enveloping by the cell 1 plasma membrane as well as an ‘exclusion layer’, likely composed of actin. II. At the onset of invasion into the second cell in the sequence (cell 2), the structures surrounding the hypha in cell 1, i.e. cell 1 plasma membrane (added during cell 1 entry), an actin layer and another cell 1 plasma membrane (added during cell 1 exit), are enveloped by the plasma membrane of cell 2. III. Invasion into cell 3 proceeds in a similar manner, with the structures surrounding the hypha in cell 2 enveloped by the plasma membrane of cell 3. Cross-section views (derived from dotted lines) show the membrane layers surrounding the hypha cell wall (black), with actin in blue, and each membrane layer coloured according to its cell of origin, i.e. magenta- cell 1, green- cell 2, yellow- cell 3. The presence of an actin layer extending within the trans-cellular tunnel may provide support to the TCT structure.

### Tunnel ‘Inflation’ lumens are directly linked to host glycogen granule stores

In accordance with our live-cell imaging experiments, we observed local ‘inflations’ of host membranes surrounding invading hyphae in three Caco-2 invasion events acquired by SBF-SEM (see **Figure 5Bi and 5C**). The inflation lumen contained multiple vesicles with a diameter of around 200 nm approximately matching that described for *C*.*a* hyphal extracellular vesicles (HEVs), though the origin of these vesicles could not be determined (Martinez-Lopez *et al*., 2020). Strikingly, in all three invasion events, host glycogen stores (previously identified by EM in Caco-2 (Biazik *et al*., 2010) in proximity to the inflation appeared to be disrupted (compared to stores not in the vicinity of inflations) with some glycogen granules found throughout what appeared to be an opening in the tunnel membranes and within the inflation lumen itself (see **Figure 5D, Movie 13**). In the data set presented here, a portion of the tunnel inflation was found outside of the host cell, though it was directly linked through a narrow passage to the intracellular inflation. The contact points between tunnel inflations and host glycogen stores may represent points of nutrient uptake from the host by *C*.*a* (see **Discussion**).

## Discussion

Our analysis of invasion scenarios in HeLa showed that both damaging and non-damaging invasion is possible in this cell line, while in Caco-2, invasion is always non-damaging. Using CM to stain host membranes enveloping invading hyphae, we show that non-damaging invasion in both cell types is always characterized by hyphae traversing host cells within trans-cellular tunnels (TCTs), in a process we term ‘*C*.*a* trans-cellular tunnelling’ (CaTCT). TCTs are also clearly observed in two of the damaging scenarios in HeLa: ‘exit site’ and ‘cell death-associated’ at least until the point of membrane breaching or host cell death, indicating that in these scenarios tunnelling also takes place, though it eventually results in damage. In the rest of HeLa invasion scenarios, continuous TCTs were either not observed (‘multiple sites’ scenario) or could not be clearly discerned (‘entry site’ scenario), suggesting that some HeLa invasion scenarios may not be associated with TCTs, though further research is required to address this possibility. Why some invasion events in HeLa result in CaTCT, while others result in damage remains an open question.

Invasion in both HeLa and Caco-2 was not altered by treatment with Cytochalasin D, a drug known to inhibit invasion via induced endocytosis. Thus, in our models, CaTCT as well as damaging invasion in HeLa is distinct from invasion by induced endocytosis. This result is consistent with previous studies suggesting that induced endocytosis does not play a role in Caco-2 invasion, though it has been suggested to occur in HeLa via an E-cadherin-independent mechanism (Dalle *et al*., 2010; Wächtler *et al*., 2012). Whether or not CaTCT is simply active penetration remains an open question. While the current view holds that only two mechanisms (induced endocytosis and active penetration) drive *C*.*a* intracellular epithelial invasion, our work, performed at the single cell level, has revealed several invasive lifestyles (CaTCT, damaging invasion associated with TCTs, and damaging invasion that may not be associated with TCTs) that may be manifestations of a single invasion mechanism, or alternatively, may be mechanistically distinct from one another at the molecular level. Addressing this question requires a systematic investigation of the fungal and host molecular mechanisms that underly CaTCT and damaging invasion scenarios. Further work is also required for understanding the interface between these lifestyles and induced endocytosis, performed in models where induced endocytosis is known to occur, such as oral and vaginal epithelial cell lines or endothelial cell lines. Ultimately, the role of CaTCT in *C*.*a* infection should be examined in more physiologically relevant models (Julian R Naglik *et al*., 2008).

CaTCT appears to be a complex process, involving significant re-organization of host membranes and compartments during hyphal extension which occurs without host membrane breaching, even as hyphae extended between invaded cells. Based strictly on structural considerations derived from fluorescence microscopy and SBF-SEM, we hypothesize that CaTCT proceeds through the following sequence of events (see **Figure 6)**: invasion into the first host cell begins with a tight enveloping of the host plasma membrane around the extending hypha, which in turn is enveloped by an actin rich layer likely derived from the host’s apical actin cortex (Schreider *et al*., 2002). These two layers constitute the TCT during invasion through the first host cell in the invasion sequence, with the actin rich layer possibly providing structural support allowing extensive host plasma membrane stretching (Drab *et al*., 2019; Kessels and Qualmann, 2021) (see **Figure 6I**). Exiting the first cell and entering the second cell in the invasion sequence adds two more membranes to the TCT structure, the first derived from the first cell’s plasma membrane upon hypha exit, and the second from the second cell’s plasma membrane upon hypha entry (see **Figure 6II**). Thus, within the second invaded host cell the TCT is composed of three membrane layers (two from the first invaded cell and one from the second invaded cell), perhaps supported by remnants of the actin rich layer found between the two membranes derived from the first invaded cell. This sequence is repeated during invasion into the third cell in the invasion sequence, with the TCT composed of five membrane layers during hyphal extension within the third cell (see **Figure 6III**). Thus, CaTCT is an iterative process, with each successive invaded cell contributing membranes to the TCT, resulting in a multi-layered tunnel structure (see **Figure 6III**).

While it is certain that the plasma membrane of each invaded cell contributes to the TCT’s membrane composition, internal membranes compartments such as the ER or the Golgi may also contribute to the TCT composition, allowing it to extend beyond the physical limits imposed by plasma membrane stretching alone. Cell polarity may also play a role in the stability of TCTs, as polar Caco-2, initially invaded in our setup exclusively via their apical side, may provide excess membrane for TCT extension via microvilli unfolding, compared to non-polar HeLa (Figard and Sokac, 2014). TCT stability may also be affected by differences in the actin cortex organization between polar and non-polar cells. Identifying the precise origins, composition and function of TCT membranes and cytoskeletal elements is an important line of investigation. Overall, CaTCT stands in stark contrast to cell-to-cell spreading of actin-motile invasive bacteria like *L. monocytogenes* and *S. flexneri* that must induce host membrane breaching followed by ‘escape’ into the host cytosol in every invaded cell before transitioning to its neighbour (Dowd, Mortuza and Ireton, 2021).

We hypothesize that TCT ‘inflations’ found in direct contact with disrupted host glycogen stores, represent sites of nutrient uptake by *C*.*a* during CaTCT. *C*.*a* access to the host’s nutrient rich cytosol during CaTCT is likely highly restricted, due to hyphae being enveloped in multi-layered TCTs. However, host glycogen stores provide an ideal site for nutrient uptake that can be readily utilized by *C. albicans* (Dennerstein and Ellis, 2001). A hypothetical mechanism could involve the export of *C*.*a* Hyphal Extracellular Vesicles (HEVs) containing glycogen phosphorylase such as Gph1 out of TCTs towards glycogen stores (Urban *et al*., 2003; Gil-Bona *et al*., 2015; Martinez-Lopez *et al*., 2020). Glycogen granules taken up into the TCT lumen via an unknown mechanism, are then broken down into glucose by cell wall glycosidases such as CaGca1p and CaGca2p (Sorgo *et al*., 2013; Maicas *et al*., 2016). This in turn leads to increased osmolarity within the TCT lumen, resulting in water influx and TCT inflation. Finally, glucose uptake into *C*.*a* hyphae restores the osmotic balance leading to a return to a tightly enveloping TCT.

As CaTCT occurs without direct contact between invading hyphae and the host cytosol (with the exclusion of the inflation-glycogen store interface) and without host damage, from an immunological standpoint, CaTCT may be perceived by host cells as an entirely extracellular process. Thus, both innate immune recognition of *C*.*a* pathogen-associated molecular patterns (PAMPs) and the activation of epithelial damage-associated molecular patterns (DAMPs) may be similar to those triggered by non-invading adherent *C*.*a* hyphae (Cohen-Kedar *et al*., 2014; Altmeier *et al*., 2016; d’Enfert *et al*., 2021; Swidergall *et al*., 2021). This hypothesis is supported by a study showing that *C*.*a* infection of Caco-2 produces only low levels of both host cellular stress responses and proinflammatory cytokines during the first hours of infection (Böhringer *et al*., 2016). Overall, CaTCT may represent a ‘commensal-invasive’ lifestyle that does not damage host cells and limits host-pathogen interactions, therefore triggering only a low-level immune response.

In conclusion, this study provides a detailed view of the invasive lifestyles of *C*.*a*. in epithelial cells, entailing either host membrane breaching or non-damaging host cell traversal. The trans-cellular tunnelling mechanism described here sheds light on long standing questions regarding the precise nature of invasion by this fungal pathogen.

## Acknowledgments

We thank R. Arkowitz and M. Bassilana (iBV, Nice, France) for providing us with the *C*.*a* strains used in this study. We thank J. Enninga and P. L. Lambert (Institut Pasteur, Paris, France) for constructing and providing us with stably expressing epithelial cell lines and other resources. We thank C. D’Enfert and S. Bachellier-Bassi (Institut Pasteur, Paris, France) for support with experimental tools. We thank M. Larsen (Cimi-Paris, Paris, France) for help with statistical analysis. EM sample preparation and SBF-SEM were performed at the ImagoSeine core facility of the Institut Jacques Monod, Paris, member of IBiSA and France-BioImaging (ANR-10-INBS-04) infrastructures. This work was supported by the ATIP-Avenir program and by the Fondation pour la Recherche Médicale (FRM) fellowship (FDM202006011423) to AP.

## Author contributions

Conceptualization, A.W.; Methodology, A.W., J.L., and J.M.V.; Investigation, J.L., A.P., D.T., and R.L.B.; Formal Analysis, J.L. and A.W.; Resources, J.M.V.; Writing – Original Draft, A.W.; Writing – Review & Editing, A.W., J.L and J.M.V.; Supervision, A.W.

## Declaration of interests

The authors declare no competing interests.

## Materials and methods

### *Candida albicans* strains and growth conditions

*C*.*a* BWP17 wild-type strain (Wilson, Davis and Mitchell, 1999) or *C*.*a* BWP17 strain expressing GFPγ-CtRac1at the plasma membrane (Bassilana and Arkowitz, 2006; Zhang and Konopka, 2010) (kindly provided by Robert Arkowitz, University of Nice-Sophia Antipolis, France) were used in all assays as indicated. Strains were routinely grown on YPD liquid/agar (1% yeast extract, 2% peptone, 2% D-glucose with or without 2% agar) supplemented with 80 µg/mL of uridine at 30°C.

### Epithelial cell lines culture

A HeLa cell line stably expressing eGFP-galectin-3 (Gal-3) was generated by Patricia Latour Lambert (Institute Pasteur, Paris, France) using the pLenti6/V5 Directional TOPO Cloning Kit (Invitrogen) with the coding sequences from the pEGFP-galectin-3 plasmid (Paz *et al*., 2010; Kreibich *et al*., 2015). All experiments on HeLa were performed using the HeLa-Gal-3 cell line. Caco-2 cell line subclone C2BBe1 stably expressing eGFP-galectin-3 (Gal-3) was generated as previously described (Rey *et al*., 2020). Cell lines were kindly provided by P. Latour Lambert and Jost Enninga (Institute Pasteur, Paris, France). Caco-2 not expressing Gal-3 were purchased from Sigma (86010202). All cells were routinely cultured in Dulbecco’s modified Eagle’s medium (DMEM) (Gibco-Invitrogen) supplemented with 10% v/v fetal bovine calf serum (Gibco-Invitrogen) at 37°C, 5% CO_2_. Live-cell imaging experiments were performed in optically transparent EM buffer (120 mM NaCl, 7 mM KCl,1.8 mM CaCl_2_, 0.8 mM MgCl_2_, 5 mM glucose and 25 mM Hepes at pH 7.3) (Weiner *et al*., 2016) supplemented with 0.2 g/L of amino acids (MP Biomedicals SC Amino Acids Mix) and 80 µg/mL of uridine.

### Infection procedure

For all experiments, epithelial cells were plated in a 96 well plate suitable for microscopy (Ibidi, polymer coverslip) 2 to 3 days or 6 to 7 days before infection for HeLa or Caco-2, respectively. HeLa were infected at 90% confluency and Caco-2 were infected 24 hours after 100% confluency was reached (Grosheva *et al*., 2020). Prior to infection, the host cells were washed two times with PBS. To label the host cell plasma membrane and endocytic compartment, CellMask Deep Red Plasma Membrane Stain or CellMask Orange Plasma Membrane Stain (ThermoFisher) was used at 2.5 µg/mL working concentration. Epithelial cells were incubated in the dark at 37°C for 10 minutes in EM medium, and washed one time with PBS. For cytochalasin inhibition experiments, host cells were treated with 0.5 µM of cytochalasin D at 37°C for 45 minutes in EM medium, and washed one time with PBS. *C*.*a* strains were cultured overnight in liquid YPD medium supplemented with 80 µg/mL of uridine at 30°C, 200 rpm shaking, followed by a back dilution at 1:100 into fresh YPD medium supplemented with 80 µg/mL of uridine and then grown to exponential phase at 30°C, 200 rpm shaking. To prevent yeast cells from clumping together, the culture was subjected to mild sonication on ice for 6 cycles of 15 seconds on and 30 seconds off (Bor *et al*., 2016), at power setting 15% using the Fisherbrand Model 50 Sonic Dismembrator (Fisher Scientific) with a standard 1/8” diameter microtip. Sonicated yeast cells were counted with a hemacytometer, diluted in EM medium and added to epithelial cells at a multiplicity of infection (MOI) of 0.1, followed by 15 minutes of incubation to allow initial adherence of yeast cells to host cells. For Caco-2 experiments, live-cell imaging was started 2 hours post-infection to maximize imaging of invasion events.

### Live-cell imaging

Live-cell imaging was performed using a fully automated inverted wide-field AxioObserver microscope (Zeiss) equipped with Colibri 7 LED illumination (Zeiss), a Hamamatsu camera (ORCA-Flash4.0 V3), a Piezo stage (Prior Scientific, NanoScanZ 100), an environmental chamber with temperature control set to 37°C and a Definite Focus 2 device (Zeiss) used to counter focus drift. For initial investigations into HeLa, a 40X air objective was used (Plan-Neofluar NA 0.6) (see **Figure 2**). For all other live-cell imaging experiments a 40X oil immersion objective (Plan-Apochromat NA 1.4) was used. Data was acquired using ZEN 2.6 pro software (Zeiss). Movies were recorded over 7 hours (such that total time post infection at the end of the acquisition was 7 hours for HeLa and 9 hours for Caco-2). Every 10 minutes, a z-stack of 15 planes with 0.94 µm z steps (40X air objective) or a z-stack of 30 planes with 0.37 µm z steps (40X oil objective) was acquired sequentially in two fluorescent channels, with a single plane acquired in phase. Gal-3 was excited with a 475 nm LED and detected with a 525 nm filter. The CellMask Orange Plasma Membrane Stain was excited with a 555 nm LED and detected with a 605 nm filter and the CellMask Deep Red Plasma Membrane Stain was excited with a 630 nm LED and detected with a 690 nm filter. The BFC stage was defined as t = 0 min for every invasion event presented in the figures.

### Differential staining invasion assay

HeLa expressing Gal-3 or Caco-2 not expressing Gal-3 were fixed with 4% PFA for 10 minutes at room temperature 4 hours or 6 hours post-infection, respectively. Infected Hela were washed 3 times with PBS and the fungal cells were firstly labelled with 10 µg/mL of concanavalin-A coupled with tetramethylrhodamine (Invitrogen) during 20 minutes at room temperature, hence labelling non-invasive fungal cells and the external parts of invasive fungal cells as previously described (Wächtler *et al*., 2012). For fixed Caco-2, a rabbit anti-*C*.*a* polyclonal antibody at 25 µg/mL (OriGene) counterstained with a secondary goat anti-rabbit IgG conjugated with Alexa Fluor 555 at 0.4 µg/mL (Invitrogen) was used as a first label as previously described (Dalle *et al*., 2010). After rinsing 3 times with PBS, the fungal cells were labelled with 25 µg/mL of Remel BactiDrop Calcofluor White (ThermoFisher) during 20 minutes at room temperature, labelling non-invasive fungal cells and both external and internal parts of invasive fungal cells as previously described (Dalle *et al*., 2010; Wächtler *et al*., 2012). After rinsing 3 times with PBS, the fixed samples were directly imaged under an inverted wide-field microscope AxioObserver (Zeiss) with a 63X oil objective (Plan-Apochromat NA 1.4). Calcofluor white was excited with a 385 nm LED and the signal was detected with a 480 nm filter. Concanavalin-A conjugated with tetramethylrhodamine or the secondary antibody was excited with a 555 nm LED and the signal was detected with a 605 nm filter. Depending on the infection site, a z-stack (mean total range of 14 µm) with 0.25 µm z steps was acquired in four channels, including phase and three fluorescent channels (Gal-3, first label, second label).

### Serial block face-scanning electron microscopy

For Serial Block Face-Scanning Electron Microscopy (SBF-SEM), Caco-2 (not expressing Gal-3) were infected by *C*.*a* BWP17 wild-type strain for 6 hours at MOI 2, then fixed in 3% PFA, 1% glutaraldehyde during 1 hour at room temperature. To increase conductivity of the sample in the microscope, the cell monolayer was covered with a thin layer of 10% gelatin containing 1% of Bovine Serum Albumine (BSA). Samples were then prepared for SBF-SEM (NCMIR protocol (Deerinck *et al*., 2010)) as follows: cells were post-fixed for 1 hour in a reduced osmium solution containing 1% osmium tetroxide, 1.5% potassium ferrocyanide in PBS, followed by incubation with a 1% thiocarbohydrazide in water for 20 minutes. Subsequently, samples were stained with 2% OsO_4_ in water for 30 minutes, followed by 1% aqueous uranyl acetate at 4 °C overnight. Cell monolayers were then subjected to *en bloc* Walton’s lead aspartate staining (Walton, 1979), and placed in a 60 °C oven for 30 minutes. Samples were then dehydrated in graded concentrations of ethanol for 10 minutes in each step. The samples were infiltrated with 30% agar low viscosity resin (Agar Scientific Ltd, UK) in ethanol, for 1 hour, 50% resin for 2 hours and 100% resin overnight. The resin was then changed and the cells were further incubated during 3 hours, prior to inclusion in upside down capsules and polymerization for 18 hours at 60 °C. The polymerized blocks were mounted onto aluminum stubs for SBF imaging (FEI Microtome 8 mm SEM Stub, Agar Scientific), with two-part conduction silver epoxy kit (EMS, 12642-14). For imaging, samples on aluminum stubs were trimmed using an ultramicrotome and inserted into a TeneoVS SEM (ThermoFisher). Acquisitions were performed with a beam energy of 3 kV, 200 pA current, in LowVac mode at 40 Pa, a dwell time of 1 µs per pixel at 10 nm pixel size. Sections of 100 nm were serially cut between images. Overall, 11 different data sets were acquired, containing 10 invasion events with hyphae entirely within the acquisition volume, and 30 invasion events containing hyphae partially within the acquisition volume.

### Image processing and analysis

Data acquired by fluorescent microscopy were analysed using ZEN 2.6 pro software (Zeiss, blue edition) and Fiji (Schindelin *et al*., 2012). Data acquired by SBF-SEM were processed using Fiji and Amira (ThermoFisher). Data alignment and manual segmentation were performed using Amira.

### Quantifications and statistical analysis

For HeLa invasion studies, a total number of 629 invasion events were recorded out of six independent replicates. The Chi-square (χ2) test was used to assess the homogeneity in the size of distribution of invasion scenarios among the six independent replicates, with a p-value threshold of 0.05 (p-value<0.0001) were recorded. For Caco-2 invasion studies, a total number of 153 invasion events were recorded in three independent experiments. To quantify the host cell death timing, cell death was assigned to an invasion stage if it occurred within 30 minutes (3 frames) of that stage’s onset. We considered host cells alive at the end of invasion if no host cell death was observed at least 90 minutes after the SC stage. Host cells that were alive after SC but were not recorded for an additional 90 min due to the end of the live imaging session were discarded for the purpose of cell death quantification. The cell traversal (CT) time was defined as the time needed for a hypha to extend through one host cell, which was calculated for every invasion event as follows: CT time = time of SC – time of FC. The CT time was calculated only for hyphae that reached the SC stage without host cell death occurring. To compare the CT time between each of the five invasion scenarios on HeLa, statistical differences were assessed with a parametric statistical test, a one-way ANOVA test (multiple t-Student test comparison) further corrected with a Tukey’s multiple comparisons with a p-value threshold of 0.05 (see **Figure S1**). For cytochalasin D treatment experiments, an identical experimental pipeline as described above was used in three independent experiments. For infected HeLa, a total number of 618 invasion events were counted out of three independent replicates (n = 311 for untreated cells and n = 307 for treated cells). The proportion of each invasion scenario was compared between treated and untreated cells with a non-parametric statistical test, a Mann-Whitney U-test, with a p-value threshold of 0.05 (p-value = 0.2 for ‘Entry site’, p-value = 0.6857 for ‘Multiple sites’, p-value = 0.3428 for ‘Exit site’, p-value = 0.8857 for ‘Cell death associated’, p-value = 0.6857 for ‘No Gal-3 recruitment’). For infected Caco-2, a total number of 261 invasion events were counted out of three independent replicates (n = 126 for untreated cells and n = 135 for treated cells). As we identified only a single invasion scenario, the number of invasion events in treated cells was normalized by the number of invasion events in untreated cells, and was compared using a non-parametric statistical test, Wilcoxon signed rank test, with a p-value threshold of 0.05 (p-value = 0.75). The difference in HeLa invasion scenario distribution in cytochalasin D treated vs. untreated cells was evaluated using a χ2 test for homogeneity with a p-value threshold of 0.05. Errors are given as standard deviation if not mentioned otherwise. Statistical analysis was performed using GraphPad Prism 9.1.2.

## Supplemental figures titles and legends

**Figure S1. Cell traversal (CT) times in HeLa and Caco-2 cells in each invasion scenario**. For HeLa cells (A), n = 414 invasion events in six independent experiments (n = 5 “Entry site”, n = 45 “Multiple sites”, n = 97 “Exit site”, n = 87 “Cell death associated”, n = 183 “No Gal-3 recruitment”) compared statistically with a one-way ANOVA test (multiple t-Student test comparison) further corrected with a Tukey’s multiple comparisons, p-value < 0.05 (p-value = 0.0706, non-significant (ns)). For Caco-2 cells (B), n = 94.

**Figure S2. Differential invasion assay in HeLa and Caco-2 cells**. HeLa or Caco-2 infections were fixed at 4 hours and 6 hours post-infection respectively. The culture was stained with Calcofluor White (CFW), labelling entire hyphae and germ cells and concanavalin-A (ConA) or an anti-*C. albicans* antibody (Ab), labelling only the non-internalized part of hyphae. Scale bars are 20 µm.

**Figure S3. High-resolution live cell imaging of *C. albicans* hyphae invading HeLa cells via the ‘entry site’, ‘cell death-associated’ and ‘no Gal-3 recruitment’ scenarios**. The invasion stages are presented as follows: before first contact (BFC); first contact with the host plasma membrane (FC); extension within the host cell (EH); second contact with the host plasma membrane (SC); after second contact (ASC). Time 0 is set to BFC. For each time point a phase image is presented together with a maximum intensity projection of three z-sections in the Gal-3 and CellMask (CM) channels, and a composite image showing Gal-3 (green) and CM (magenta) together. A cell death event is marked with a red ‘x’. Cross-sections derived from the dotted line in the corresponding time point are presented in the Gal-3 and CM channels. Scale bars are 10 µm, and 5 µm in the cross-section views.

**Figure S4. Single SBF-SEM sections (in the XY plane) showing hyphae from three different Caco-2 invasion events**. Several host membrane layers surrounding each hypha are observed. Scale bars are 1 µm.

## Supplemental movies titles and legends

**Movie 1, corresponding to figure 1B -** HeLa invasion, **‘entry site’ scenario**. From left to right: phase, Gal-3, CM and a Gal-3 (green) and CM (magenta) composite view are presented. Time points match those presented in figure 1B.

**Movie 2, corresponding to figure 1B -** HeLa invasion, **‘multiple sites’ scenario**. From left to right: phase, Gal-3, CM and a Gal-3 (green) and CM (magenta) composite view are presented. Time points match those presented in figure 1B.

**Movie 3, corresponding to figure 1B -** HeLa invasion, **‘exit site’ scenario**. From left to right: phase, Gal-3, CM and a Gal-3 (green) and CM (magenta) composite view are presented. Time points match those presented in figure 1B.

**Movie 4, corresponding to figure 1B -** HeLa invasion, **‘cell death-associated’ scenario**. From left to right: phase, Gal-3, CM and a Gal-3 (green) and CM (magenta) composite view are presented. Time points match those presented in figure 1B.

**Movie 5, corresponding to figure 1B -** HeLa invasion, **‘no Gal-3 recruitment’ scenario**. From left to right: phase, Gal-3, CM and a Gal-3 (green) and CM (magenta) composite view are presented. Time points match those presented in figure 1B.

**Movie 6, corresponding to figure 2 -** HeLa invasion, high resolution. From left to right: phase, Gal-3, CM and a Gal-3 (green) and CM (magenta) composite view are presented. Time points match those presented in figure 2.

**Movie 7, corresponding to figure 3A -** Caco-2 invasion scenario. Top left: phase, top right: Gal-3, bottom left: CM, bottom right: Gal-3 (green) and CM (magenta) composite view. Time points match those presented in figure 3A.

**Movie 8, corresponding to figure 3B -** 3D segmentation of CM labelling during Caco-2 invasion. CM labelling within the entire acquisition volume is segmented in three dimensions (magenta). Background (orange) provides host cells outline. Time points match those presented in figure 3B, for both XY and XZ views.

**Movie 9, corresponding to figure 3D -** ‘Inflation’ during Caco-2 invasion. From left to right: phase, *C*.*a*, CM and a *C*.*a* (green) and CM (magenta) composite view are presented. Time points match those presented in figure 3D.

**Movie 10, corresponding to figure 5A -** SBF-SEM, overview of invasion site. Data binned to a XY resolution of 20 nm is presented. Axes match those in figure 5A. Slices from the data set are presented in sequence, leading to the apparent “movement” in the movie. Segmentations are as follows: white- *C*.*a* hypha, magenta- host cell 1, green- host cell 2, yellow- host cell 3, brown- host cell nuclei and blue- host glycogen stores.

**Movie 11, corresponding to figure 5B -** SBF-SEM, The organization of host membranes around the hypha in each invaded cell. Data binned to a XY resolution of 20 nm is presented. Segmentation is the same as in movie 10.

**Movie 12, corresponding to figure 5C -** Cross-section view of trans-cellular tunnel. Data binned to a XY resolution of 20 nm is presented via a non-orthogonal slice (in relation to the acquisition volume). Axes match those in figure 5C. Hypha is segmented in white. Trans-cellular tunnel segments are segmented according to their association with host cells the invasion sequence: magenta- host cell 1, green- host cell 2, yellow- host cell 3.

**Movie 13, corresponding to figure 5D -** Direct contact between ‘inflation’ lumen and host glycogen granules. Data binned to a XY resolution of 20 nm is presented. Segmentation matches that in movie 12 with the addition of host glycogen store in blue. The *C*.*a* Spitzenkörper is also observed.

## Notes

### Competing Interest Statement

The authors have declared no competing interest.

