## Supplemental figures for "Trans-cellular tunnels induced by the fungal pathogen *Candida albicans* facilitate invasion through successive epithelial cells without host damage"

**Supplemental Data**

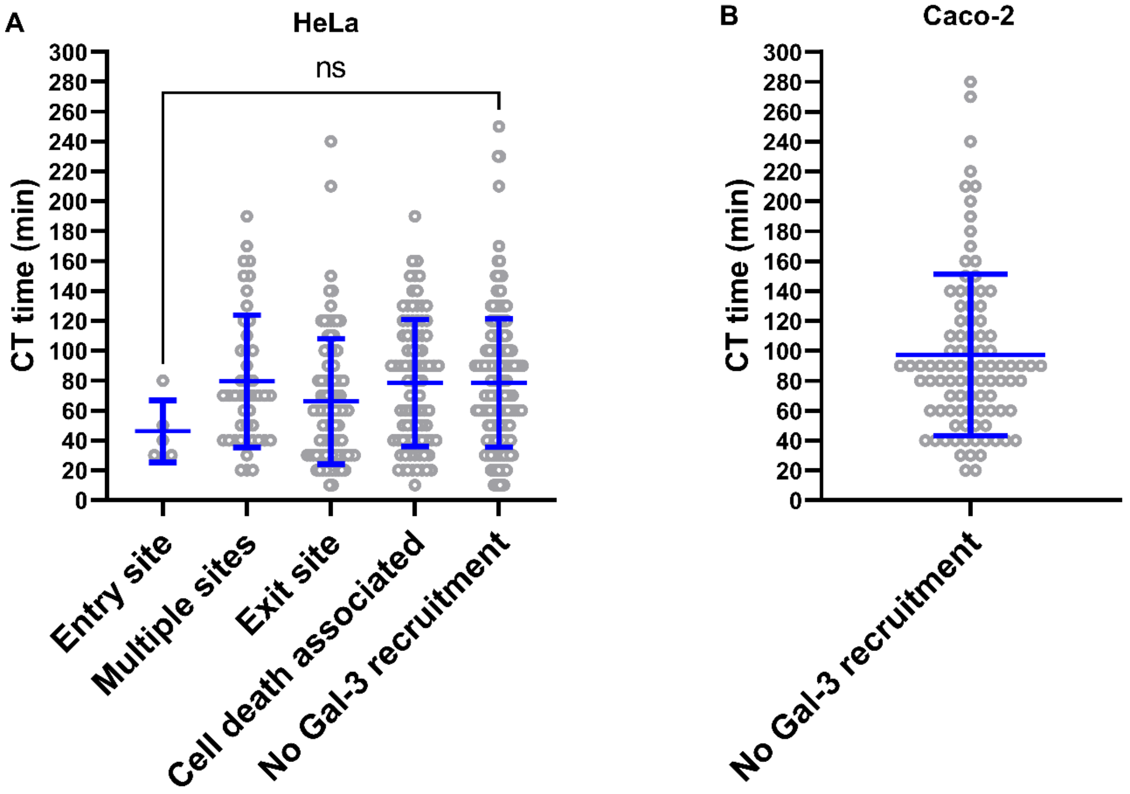


**Figure S1**. **Cell traversal (CT) times in HeLa and Caco-2 cells in each invasion scenario**. For HeLa cells (A), n = 414 invasion events in six independent experiments (n = 5 “Entry site”, n = 45 “Multiple sites”, n = 97 “Exit site”, n = 87 “Cell death associated”, n = 183 “No Gal-3 recruitment”) compared statistically with a one-way ANOVA test (multiple t-Student test comparison) further corrected with a Tukey’s multiple comparisons, p-value < 0.05 (p-value = 0.0706, non-significant (ns)). For Caco-2 cells (B), n = 94.

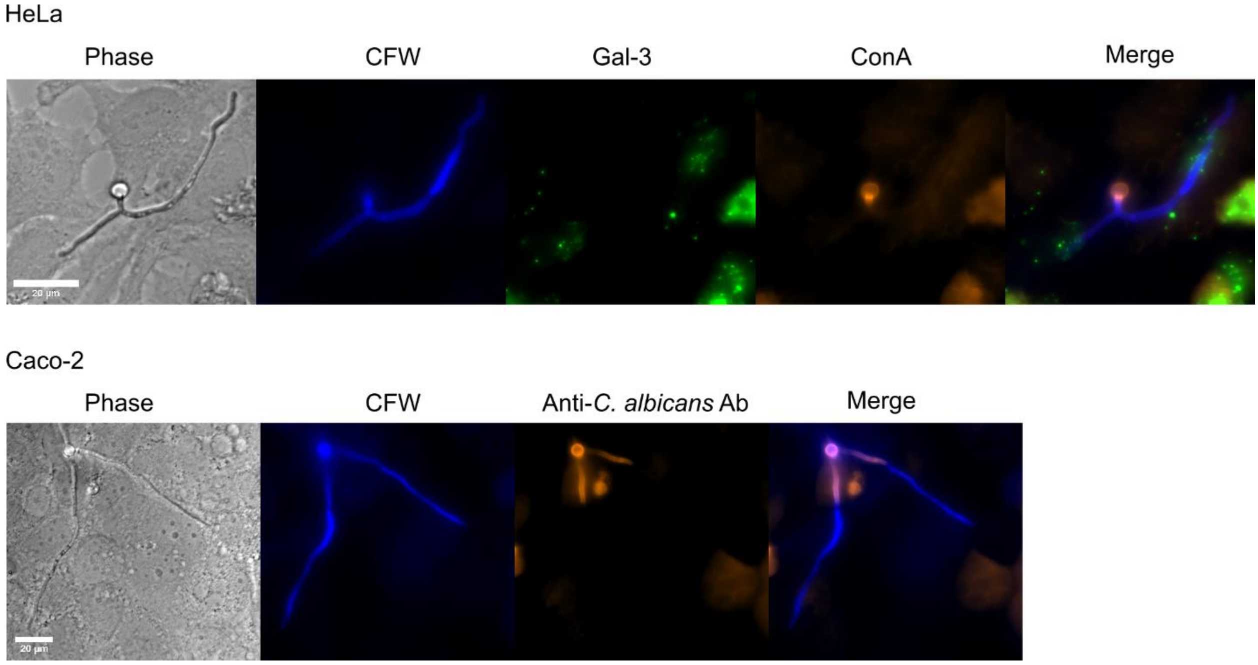


**Figure S2. Differential invasion assay in HeLa and Caco-2 cells.** HeLa or Caco-2 infections were fixed at 4 hours and 6 hours post-infection respectively. The culture was stained with Calcofluor White (CFW), labelling entire hyphae and germ cells and concanavalin-A (ConA) or an anti-*C. albicans* antibody (Ab), labelling only the non-internalized part of hyphae. Scale bars are 20 µm.

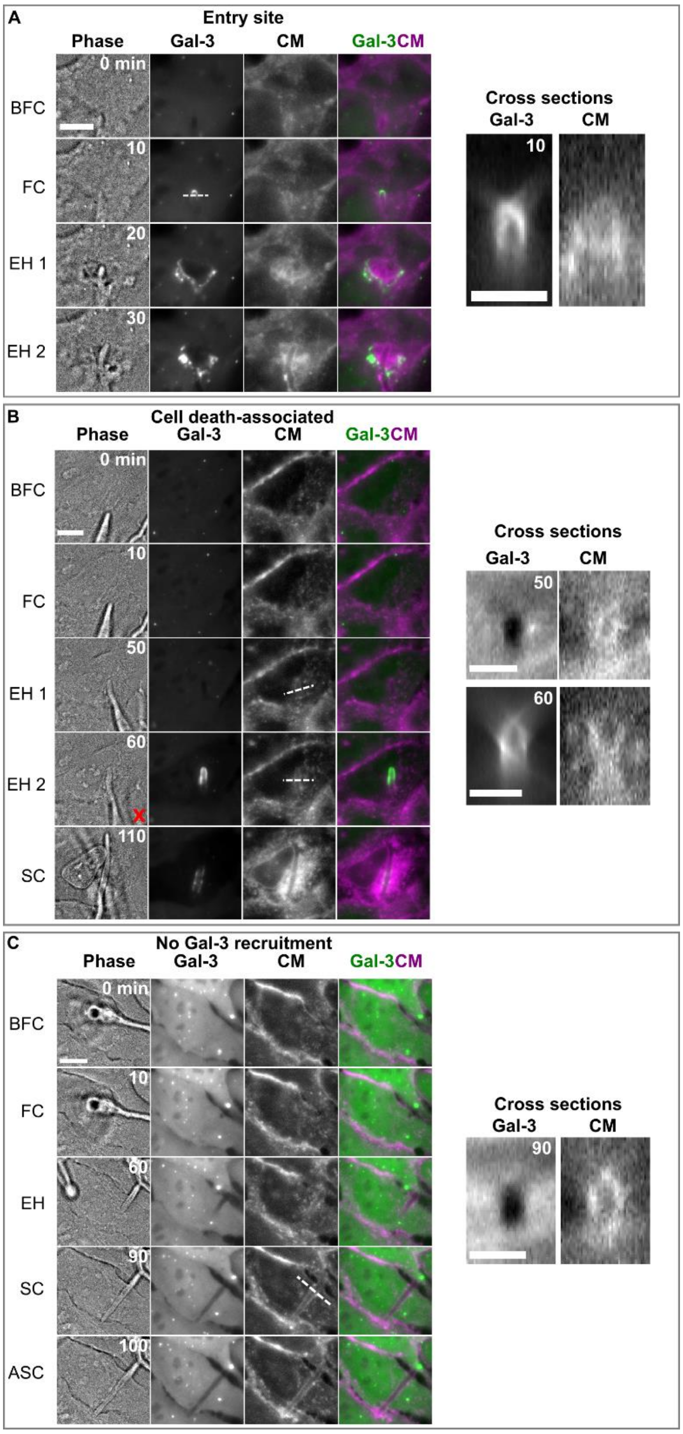


**Figure S3. High-resolution live cell imaging of *C. albicans* hyphae invading HeLa cells via the 'entry site', 'cell death-associated' and 'no Gal-3 recruitment' scenarios.** The invasion stages are presented as follows: before first contact (BFC); first contact with the host plasma membrane (FC); extension within the host cell (EH); second contact with the host plasma membrane (SC); after second contact (ASC). Time 0 is set to BFC. For each time point a phase image is presented together with a maximum intensity projection of three z-sections in the Gal-3 and CellMask (CM) channels, and a composite image showing Gal-3 (green) and CM (magenta) together. A cell death event is marked with a red ‘x’. Cross-sections derived from the dotted line in the corresponding time point are presented in the Gal-3 and CM channels. Scale bars are 10 µm, and 5 µm in the cross-section views.

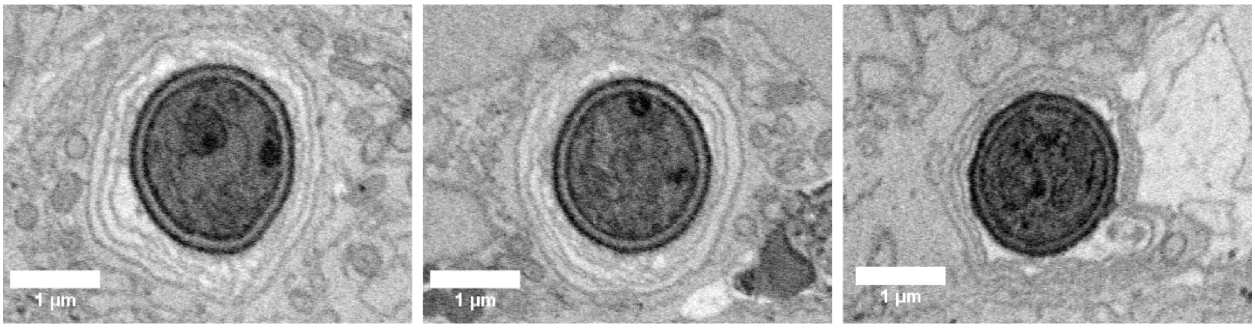


**Figure S4**. **Single SBF-SEM sections (in the XY plane) showing hyphae from three different Caco-2 invasion events**. Several host membrane layers surrounding each hypha are observed. Scale bars are 1 µm.
